# Meta-Analysis of the Functional Neuroimaging Literature with Probabilistic Logic Programming

**DOI:** 10.1101/2022.02.25.481931

**Authors:** Majd Abdallah, Valentin Iovene, Gaston Zanitti, Demian Wassermann

## Abstract

Inferring reliable brain-behavior associations requires synthesizing evidence from thousands of functional neuroimaging studies through meta-analysis. However, existing meta-analysis tools are limited to investigating simple neuroscience concepts and expressing a restricted range of questions. Here, we expand the scope of neuroimaging meta-analysis by designing NeuroLang: a domain-specific language to express and test hypotheses using probabilistic first-order logic programming. By leveraging formalisms found at the crossroads of artificial intelligence and knowledge representation, NeuroLang provides the expressivity to address a larger repertoire of hypotheses in a meta-analysis, while seamlessly modelling the uncertainty inherent to neuroimaging data. We demonstrate the language’s capabilities in conducting comprehensive neuroimaging meta-analysis through use-case examples that address questions of structure-function associations. Specifically, we infer the specific functional roles of three canonical brain networks, support the role of the visual word-form area in visuospatial attention, and investigate the heterogeneous organization of the fronto-parietal control network.

## 1 Introduction

Meta-analysis confronts hypotheses of brain-behavior mappings with synthesized evidence from thousands of functional neuroimaging studies [1]. However, commonly used standard tools for meta-analysis, such as BrainMap [2] and Neurosynth [3] are limited in their *formal expressivity*. For example, it is challenging to express complex questions like ‘which topics are likely to be present in a study given activation in one set of regions and *there exists no* activation in another set of regions’. Answering this question requires that we explicitly run a meta-analysis on every single term of interest, while integrating heterogeneous and uncertain data, such as spatial masks and parcellations, to define the regions [4–7]. Importantly, however, applying the nontrivial criteria to select articles for inclusion and exclusion necessitates coding in a general-purpose programming language, which can be error-prone and hard to maintain. In this work, we harness advancements in symbolic artificial intelligence, specifically in *probabilistic logic programming*, to design *NeuroLang* : a domain-specific language (DSL) to formulate rich and expressive neuroscience hypotheses using formal logic-based criteria, while seamlessly combining heterogeneous data and reasoning about uncertainty. Our bet is that NeuroLang’s clear syntax and probabilistic logic semantics will enable accessible, rigorous, and highly reproducible large-scale meta-analysis.

Large-scale meta-analysis has become popular in cognitive neuroscience research as a result of the growth in non-invasive neuroimaging experiments, which provoked an upsurge in the number of yearly publications on structure-function associations [8]. As a powerful approach to synthesizing large quantities of results, meta-analysis has been performed to distinguish spurious from replicable findings, derive coactivation patterns [9], and define a priori regions of interest [10]. Moreover, elaborate coactivation-based connectivity, meta-analytic functional parcellations, and sophisticated network models are now being inferred from datasets covering tens of thousands of subjects. Recently, even more elaborate meta-analyses have been performed to derive new findings. For instance, Yeo et al. [6] have used meta-analysis to study the human association cortex, identifying frontal and parietal regions that are either specialized or flexible. Furthermore, meta-analysis has been used to support findings on the fractionation of networks into subsystems underlying disparate processing domains [7, 11]. Thus, large-scale metaanalysis is essential for the continued detection of latent properties of brain systems, far beyond what can be inferred from individual studies.

Over three decades ago, spatial normalization—the use of peak activation coordinates standardized in stereotaxic space—was introduced to human neuroimaging research [12]. This norm has been embraced, enlarged, and popularized by an outstanding series of methodological breakthroughs in the field. As a result, a large and methodologically cohesive functional neuroimaging literature has emerged over the past years. Capitalizing on this wealth of results, large-scale online databases have been created to compile peak activation coordinates and their related meta-data, spurring the development of coordinate-based meta-analytic approaches to exploit this rapidly growing corpus. This makes coordinate-based meta-analysis (CBMA) the most popular type of neuroimaging meta-analysis so far. The earliest approach to compiling peak coordinates, used by BrainMap [2], is to manually transcribe them from study tables. The peaks are then linked to the content of the studies by taxonomy experts. A more recent approach, used by Neurosynth [3], is to automate this laborious task via special software that automatically extracts peak coordinates from study tables. Neurosynth also automates the annotation of studies using natural language processing techniques that incorporate term-frequency features, i.e. TF-IDF, estimated from the texts of studies. Albeit noisier, these fully automated approaches to meta-analysis scale better to the rapidly expanding neuroimaging literature.

Tools like BrainMap [2] and Neurosynth [3] have indeed simplified meta-analysis, becoming cornerstones of contemporary neuroscience research. However, only a narrow range of questions can be *natively* posed using these tools due to their limited *expressivity*. Specifically, these tools are based on propositional logic, which makes formulating queries beyond straightforward propositions, while applying nontrivial selection criteria of studies, a verbose and arduous task. More recently, NeuroQuery, a regularised predictive model trained on CBMA data, has been introduced to map *any arbitrary text fragment* (i.e. a set of keywords of interest, dubbed ‘query’) to a brain activation map. This approach shows that modelling semantic relations across studies can produce meaningful statistical maps for terms that are rarely mentioned in the literature. Nonetheless, NeuroQuery cannot express questions that infer a pattern of activation from studies associated with ‘working memory’, for instance, *but not associated* with other related functions. Such an analysis could be interesting to infer unique neural substrates among closely-related mental functions. Yet, NeuroQuery does not employ any *logic-based semantics*. That is, the words ‘but’ and ‘not’ are not in NeuroQuery’s pre-selected vocabulary, and have no influence on the resulting statistical maps. Thus, there seems to be a space for a logic-based framework that, by giving access to more elaborate meta-analytic queries, could help bridge the gap between statistical modeling and cognitive neuroscience. An all encompassing domain-specific language (DSL) for neuroscience research could encode any type of data, express complex queries using logic semantics, and reason about elements of uncertainty, all in a single unifying framework.

Parallel to the growth of functional neuroimaging over the past few decades, the artificial intelligence (AI) sub-fields of *probabilistic logic programming* and *probabilistic databases* have experienced a rapid evolution. This has lead to the development of methods that efficiently represent knowledge with both *logical* and *probabilistic* semantics in a way that makes statistical model assumptions declarative, clear and less biased. Theoretical formalisms ground probabilistic logic languages with well-defined semantics and a mathematical understanding of the complexity of various classes of queries and different types of data [13]. Efforts to address the scalability of probabilistic logic systems led to recent algorithms and data structures that can remarkably speed up analyses, even on very large and diverse databases [14].

Can probabilistic logic programming formalize and broaden the range of questions that can be expressed in a meta-analysis? Can it simplify formulating complex hypotheses while combining heterogeneous and uncertain data? In this work, we present NeuroLang, a domain-specific language for conducting comprehensive neuroimaging meta-analyses using formal, succinct, and self-contained programs. At its core, NeuroLang uses predicate logic, as opposed to propositional logic, to ease the formulation of queries in a way that is closer to human discourse [15], and that can be run against structured neuroscientific data. Logic rules and queries are augmented with probabilistic semantics to account for uncertainty that emerges from missing information, analytical variability across studies, and measurement imperfections. In this article, we present concrete use-case applications of NeuroLang that address popular questions from the literature.

## 2 Results

The use-case examples of NeuroLang shed light on the utility of probabilistic logic semantics in representing neuroscience hypotheses that cannot be readily expressed with standard meta-analysis tools. In the first example, we use between-network segregation queries to infer the unique functional roles of three canonical functional networks: the dorsal attention network (DAN), default mode network (DMN), and frontoparietal cognitive control network (FPCN). In the second example, we explore potential associations between topics and activity within the visual word-form area (VWFA), when it either coactivates with regions of the dorsal attention network or those of the language network. In the third and fourth examples, we study the functional heterogeneity of the FPCN, uncovering differential activation profiles for a number of mental functions and varying connectivity patterns with other brain networks.

### Representing Neuroscientific Knowledge under Uncertainty

Before exploring use-case examples of NeuroLang, we describe how heterogeneous neuroimaging data are represented in using fact and rule *tables*. A table is a set of tuples or rows, each representing a data instance and has a set of *k* elements representing columns. *Probabilities* can be ascribed to the rows of a table to quantify the level uncertainty in the data presented by each, in which case the table is said to be *probabilistic*.

Studies in a CBMA database report a set of *peak* activations that we store in a table named PeakReported. This table contains one row (*x, y, z, s*) for each peak that a study *s* has reported at location (*x, y, z*) in MNI space. Moreover, the uncertainty around the spatial location of peaks can be encoded in a rule table by assuming each peak’s 10mm neighboring voxels to be equivalently reported, similar to the multilevel kernel density analysis (MKDA) [16]. This rule table is called VoxelReported, and it includes a row (*x, y, z, s*) for each voxel at location (*x, y, z*) in the neighborhood (< 10*mm*) of a peak reported by study *s*. More details on how other spatial smoothing priors can be encoded in NeuroLang, such as the probabilistic prior used by the activation likelihood estimation (ALE) [17] algorithm, are provided in the Supplementary Materials.

Further, each study within a meta-analytic corpus is associated with cognitive processes or concepts addressed by its experiments. Fully automated meta-analytic tools like Neurosynth calculate statistical term-frequency features on study texts or abstracts, and threshold them to establish these links in a data-driven manner [3]. We store these associations within a TermAssociation table, containing one row (*t, s*) for each term *t* associated with study *s*. Moreover, we incorporate data-driven topic models, learned and openly shared by Neurosynth [18], within a TopicAssociation probabilistic table, containing one row (*t, s*, **P**) for each uncertain association between a topic *t* and a study *s*. In probabilistic logic, we write TopicAssociation(*t, s*) :: **P** to state ‘study *s* has a probability **P** of being associated with topic *t*‘[19]. This data representation process is illustrated in fig. 1.

**Figure 1:**
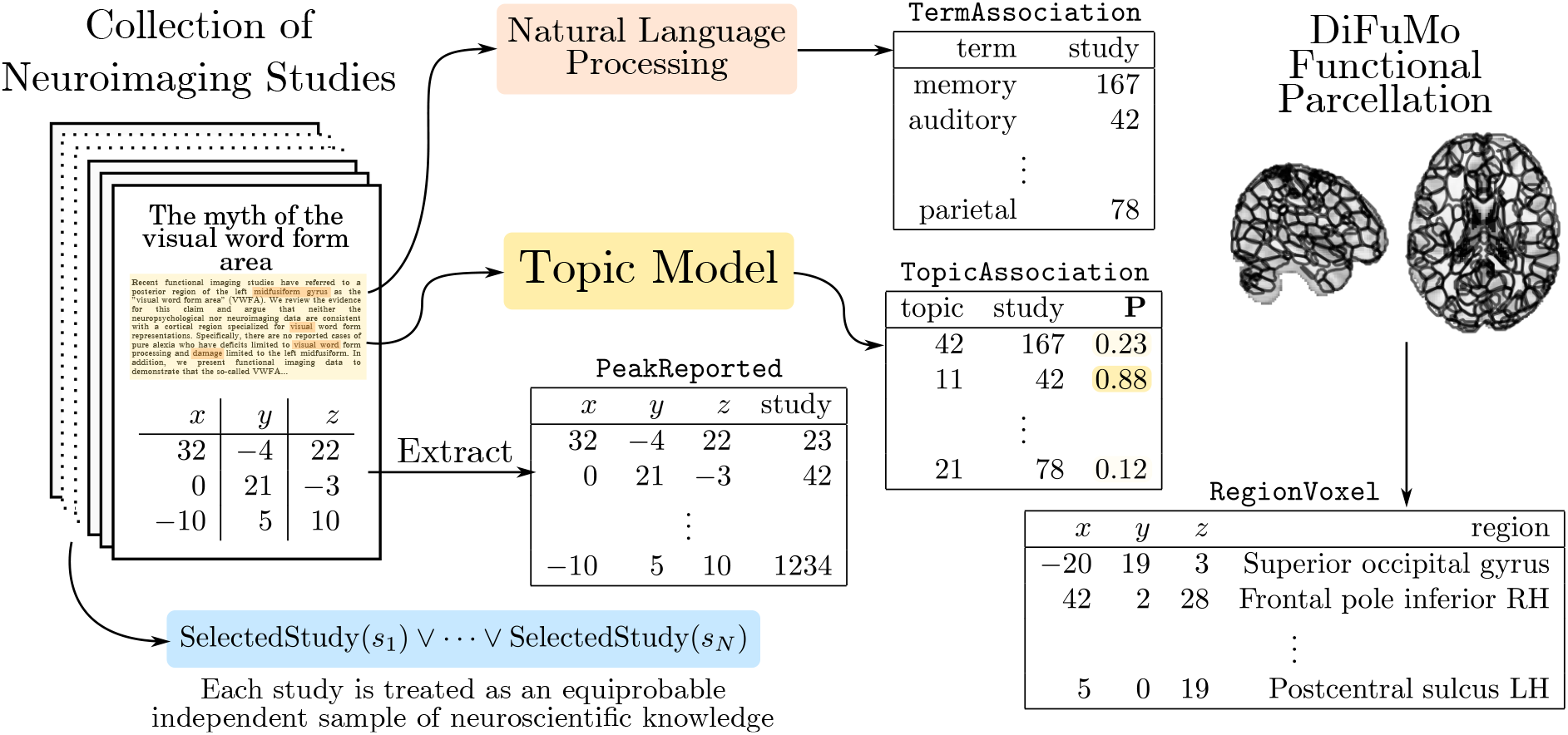
Representation of meta-analytic and functional parcellation knowledge using database tables in NeuroLang.

Similarly to Neurosynth, we assume each study within the meta-analytic database to be an *independent equiprobable sample* of neuroscientific knowledge [3, 17]. This assumption is encoded by a SelectedStudy rule, depicted in the bottom left part of fig. 1, which gives studies an equal weight 1*/N* in any meta-analysis, where *N* is the total number of studies within the meta-analytic database. This makes it possible to estimate statistics on CBMA databases in the absence of statistical power indicators (e.g. sample size).

It is common for meta-analyses to integrate anatomical or functional brain parcellations [11] to enhance interpretability and reduce computational burdens. In the examples, we use the DiFuMo-256 atlas [20], which is part of a multiscale “soft” parcellation estimated from thousands of subjects across 27 studies that include both task and resting-state fMRI experiments. This data-driven functional atlas is argued to achieve comparable statistical performance as voxel-level analyses, while simultaneously reducing computational cost and enhancing interpretability. We represent the 256 functional regions from DiFuMo in a RegionVoxel table, containing a row (*r, x, y, z*) for each brain voxel at MNI location (*x, y, z*) belonging to a DiFuMo-256 region *r*. An excerpt of the RegionVoxel table is depicted on the right part of fig. 1. We also incorporate a table NetworkRegion that contains a row (*n, r*) for each region *r* that significantly overlaps with some network *n* from the 7 or 17-network parcellations [21]. The network membership of regions is provided as part of the DiFuMo meta-data file. The experiments that follow will use this unified framework of knowledge representation to formulate probabilistic logic programs that drive meta-analytical findings.

### Between-Network Segregation: Reverse Inference of Brain Network Function

In this example, we perform a *segregation-based meta-analysis* to infer the likelihood of a topic to be present in a study given activation in a brain network, with an additional constraint that there *exists no activation in other networks*. The goal of this example is to show that a segregation query can identify which network’s activation pattern is preferentially more predictive of the presence of topic terms related to certain mental functions.

We use the Neurosynth CBMA database [3], consisting of 14,371 studies, and its associated v5-topics-100 topic model [18]. The networks included in this example are the DMN, FPCN and DAN defined using the coarse 7-Network atlas [21]. These networks exhibit coupling dynamics in support of an array of internally and externally-directed mental functions [22]. However, each one of them is believed to subserve a unique set of cognitive processes [7, 22, 23]. The FPCN contributes to a wide variety of tasks by engaging top-down control processes, the DAN is concerned with orienting attention towards salient cues, and the DMN is involved in abstract self-referential, social and affective functions. Using a segregation query, we can quantitatively identify the specific functional roles of these networks from the literature.

First, we have to represent useful heterogeneous data in NeuroLang. For instance, we assume a DiFuMo-256 component *r* to be reported by a study *s* whenever a peak activation is reported by the study within that region. In NeuroLang, this is expressed by the following logic rule

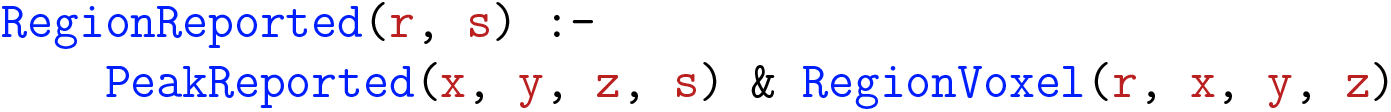

which translates, in plain English, to ‘region *r* is reported by study *s* if *s* reports a peak at location (*x, y, z*) that falls within region *r*’. Furthermore, we model the reporting of networks by studies in a *probabilistic* table. The probabilities are based on the total volume of the reported regions that belong to a network. This table accounts for the uncertainty in the location of reported peak activation coordinates as well as the number of potentially reported regions. More precisely, we consider that each study has a probability of reporting a network, proportional to the number of reported regions belonging to the network.

This is implemented by the following rules in NeuroLang

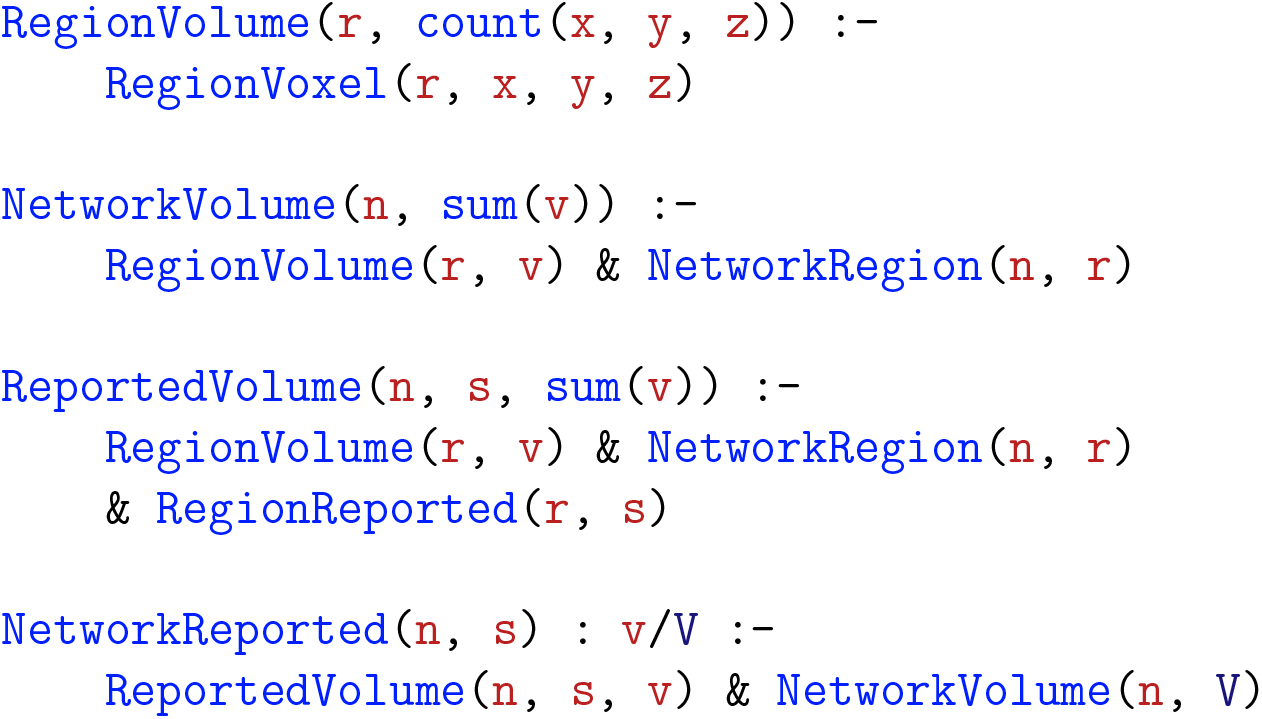

In plain English, ‘a network *n* is considered to be reported by study *s* with probability *v/V*, where *v* is the total volume of regions within network *n* that are reported active by study *s*, and *V* is the total volume of all regions in the network’. This program makes use of NeuroLang’s built-in count and sum aggregation functions.

Next, we define a rule that infers the probability that studies are associated with a topic *given* activation in only one of the three networks. That is, we query the probability that a study *s* is associated with topic *t* given that some network *n* is reported by *s* and *there exists* no other network reported by study *s*. In NeuroLang, this corresponds to the following rule

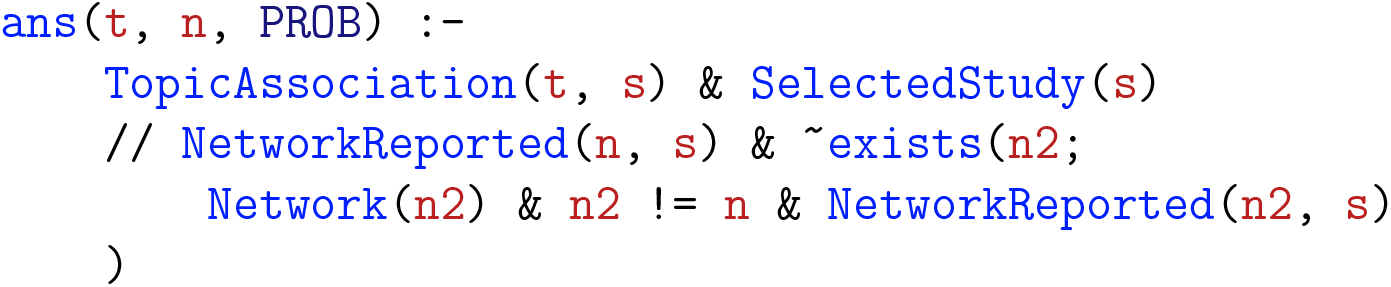

where the // operator is read as *given*, representing *probabilistic conditioning*. This rule contains a negated existential expression, ∼exists(…), that prevents two or more networks from being reported by a study at the same time.

We report the resulting functional profiles in fig. 2. We observe that topics related to sensory processing of direct environmental demands such as eye movements, visual attention, and spatial orientation are more likely to appear in studies reporting activations in the DAN only. Also, we observe that topics related to cognitive control such as task switching, task demands, response inhibition, and performance monitoring are more likely to be mentioned in studies reporting activations in the FPCN. Finally, topics related to higher-order abstract cognitive and memory-related processes are mostly associated with studies reporting DMN activations only.

**Figure 2:**
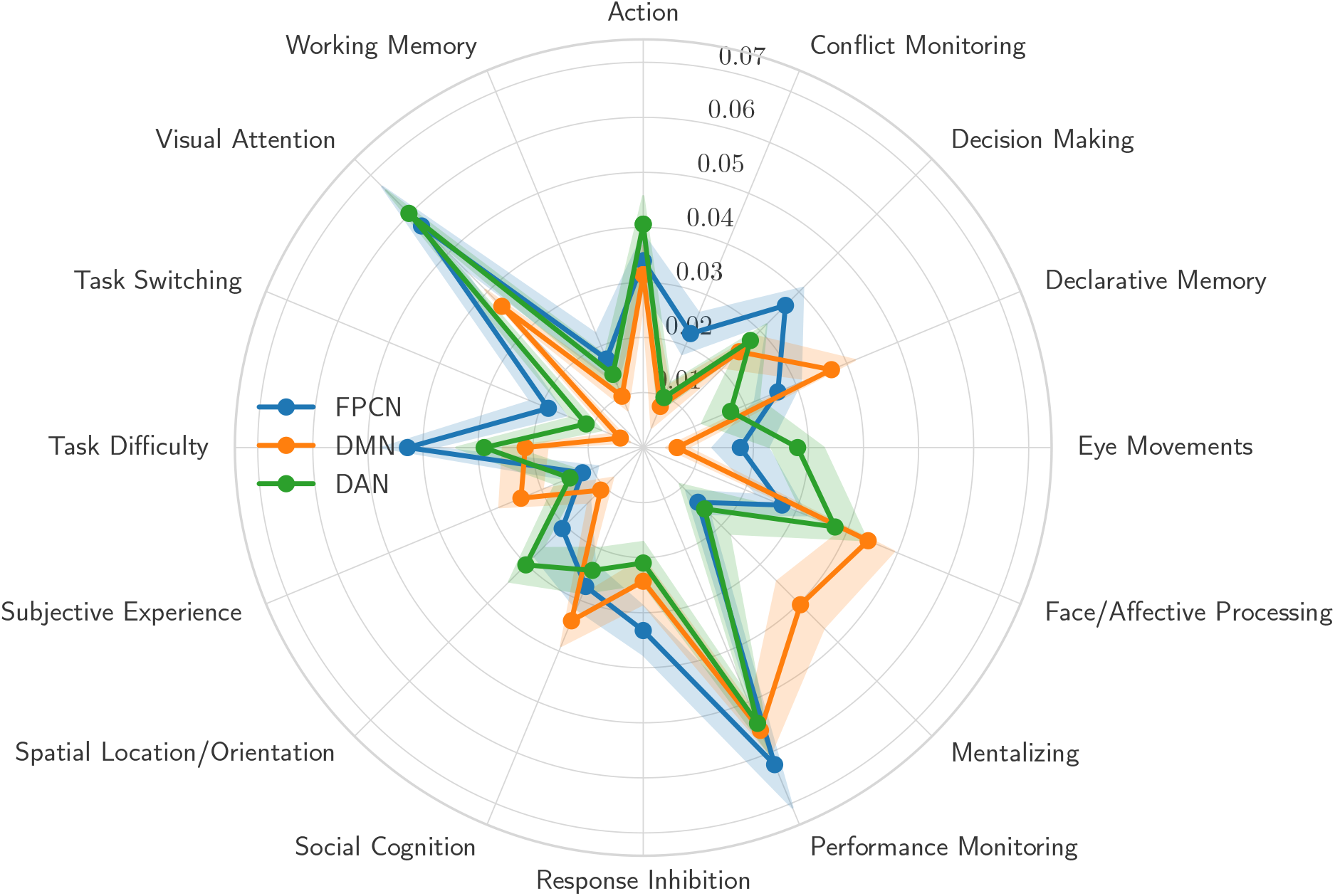
Functional profiles obtained with network-based segregation queries that identify the most probable topic associations in studies reporting activations within one network but not reporting activations within any of the other networks. A 95% confidence interval is depicted, across 1000 random 50% subsamples of the Neurosynth database [3].

### Meta-Analysing the Role of the Visual Word-Form Area in Attention Circuitry

The visual word-form area (VWFA) has attracted controversy over the years with recent findings suggesting it to take part in the attention circuitry not only in the language network [24]. Can this relationship be inferred solely from a meta-analysis of past studies that have reported activations in the left ventral occipito-temporal cortex without necessarily identifying it as the VWFA?

To answer this question, we formulate queries that infer the most probable topic associations among studies that report activations close to the VWFA region, while simultaneously reporting activations within regions of the attention network, but *not* reporting activations within regions of the language network.

To define regions of the VWFA; dorsal attention and language network, we use the seed locations defined in [24], and store them in a RegionSeedVoxel table. This table contains a row (*x, y, z, r*) for each region *r*’s seed location (*x, y, z*). A database table NetworkRegion contains rows (*n, r*) for each region *r* belonging to network *n*. A brain region is considered to be reported by a study if it reports a peak activation within 10mm of the region seed location. The choice of a 10mm radius was used to facilitate comparisons with the range of smoothing kernels that are typically used within metaanalyses.

In this example, a network is considered to be reported by a study if it reports one of the network’s regions. In NeuroLang, these rules are as follows:

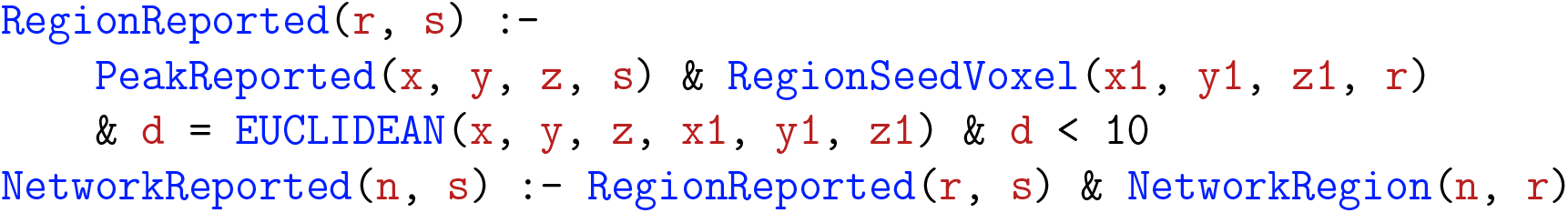

where EUCLIDEAN is a built-in function that calculates the Euclidean distance between two locations in MNI space.

Finally, to test our hypothesis, we use the following *probability encoding rule*

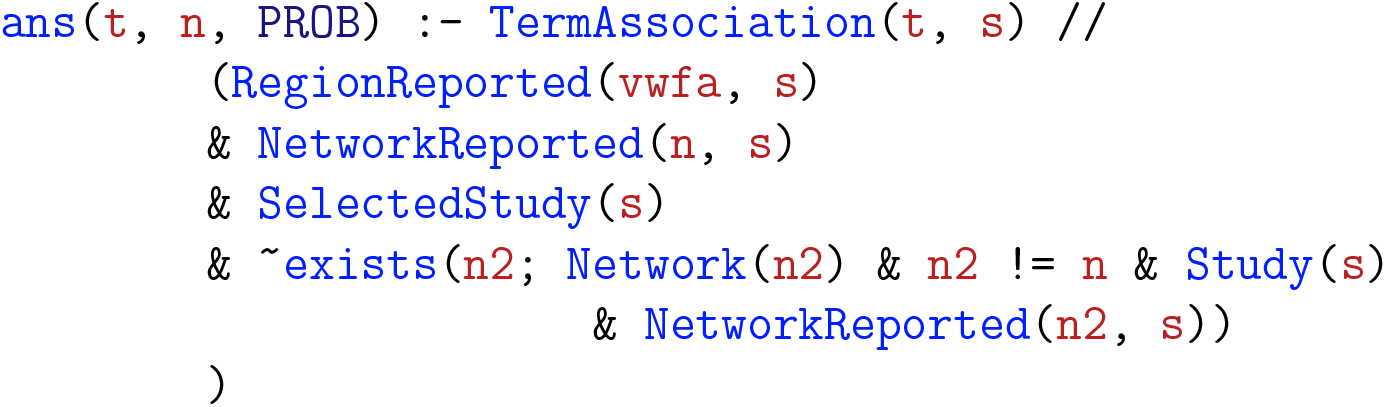

which calculates the probability of finding an association with topic *t* among studies that report the activation of both the VWFA and network *n*, but do not report the activation of any other network *n*_2_, where *n*_2_ ≠ *n*. Because only two networks, language and attention, are present in the Network table, this rule simultaneously calculates the probabilities for each pair of networks, including one while segregating the other.

Results are shown in table 1. Topic 32 was found to be significantly associated with studies that report activations within the VWFA and the attention network but that do not report activations within the ‘language’ network. This topic loads on terms related to *object recognition*—a task for which attention circuitry is essential [25]. This result suggests that the VWFA may play a role in attention, as studies that report its activations are significantly associated with object recognition, and supports the running hypothesis that the VWFA plays a role in processing multiple categories of visual stimuli [24]. We also observe a significant association with topic 21, which loads on terms related to the task of *reading words*—the putative role of the VWFA.

**Table 1:**
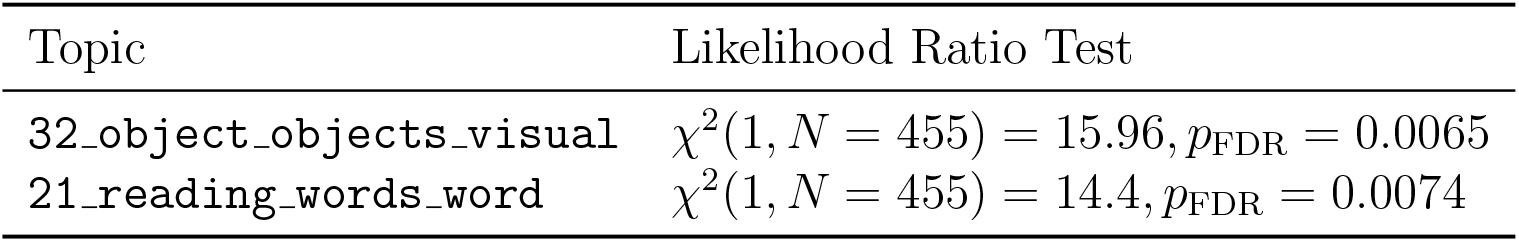
Topics associated with studies reporting the VWFA and the fronto-parietal attention network, but not reporting the ‘language’ network. We only depict associations surviving a likelihood ratio test and a false discovery rate (FDR) correction for multiple comparison (*p*_FDR_ < 0.05). More details on the likelihood ratio test are provided in the Supplementary Materials. Topics are those from Neurosynth’s v5-topics-100 topic model. Out of Neurosynth’s 14,371 studies, 455 report activations within the frontoparietal attention network without reporting activations within the ‘language’ network.

The *opposite* segregation query selects studies reporting the VWFA and the ‘language’ network but not reporting the attention network (*N* = 318). This analysis did not yield any significant topic association after correction for multiple comparison. However, a similar topic association analysis, but without segregating studies that report activation in the attention network, does yield a significant association with topic 21, linked to the ‘reading words’ (*χ*^2^(1, *N* = 852) = 56.86, *p*_FDR_ = 0.000081). This result might have more than one explanation, but a plausible explanation could be the relative decrease in statistical power (i.e. smaller number of studies) in the segregation query compared to the non-segregation query.

## Inferring Differential Activation Patterns within the FPCN using Topic Segregation Queries

In this example, we perform forward inference using *topic-based segregation queries* to derive activation patterns within the frontoparietal cognitive control network (FPCN). As a major part of the *multiple demand* system [26], the FPCN is associated with a large set of tasks, themselves belonging to disparate and overlapping cognitive processes such as working memory, memory retrieval, task switching, and semantic processing, to name a few. Moreover, there is evidence for a heterogeneous internal organization in the FPCN, whereby a different combination of regions may be involved in a different domain of control processing [27]. Thus, the goal of this example is to infer activation patterns within the FPCN predicted by the presence of topic terms related to one process and the simultaneous absence of topic terms related to other processes. In this sense, segregation queries can enhance the relative specificity of meta-analytic forward inferences by minimizing the amount of overlap amongst related topics.

From the set of 200 Neurosynth topics (version-5), we select five exemplar topics representing a subset of the cognitive processes often attributed to the FPCN, along with the loading values of studies on each topic. These topics are *working memory, decision making, task set switching, semantic control*, and *memory retrieval* [22, 26, *28, 29]. Then, we formulate the following NeuroLang program which performs topic segregation queries, yielding an activation map for each topic separately. This program is written as follows:*

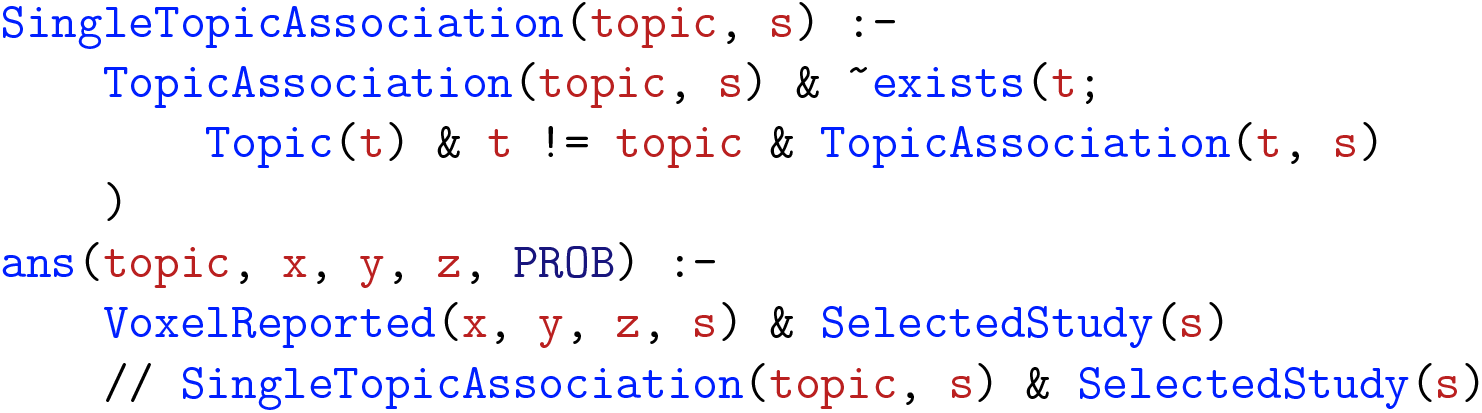

We report the resulting topic-based activations within the FPCN in fig. 3. The results of this segregation query show that the FPCN exhibits a varied activation profile across topics, corroborating previous findings of flexible adaptation of activity within this network as task demands change. Specifically, working memory and task set switching tend to activate, to some extent, spatially interleaved, frontal and parietal regions of the FPCN network. Semantic processing, on the other hand, dominantly activates a left-lateralized ventral frontal regions. Finally, decision making and memory retrieval are associated with activation in the cingulo-medial portion of the FPCN, the pre-supplementary motor/dorsal anterior cingulate cortex (decision making) and a precuneus/posterior cingulate cortex network (memory retrieval).

**Figure 3:**
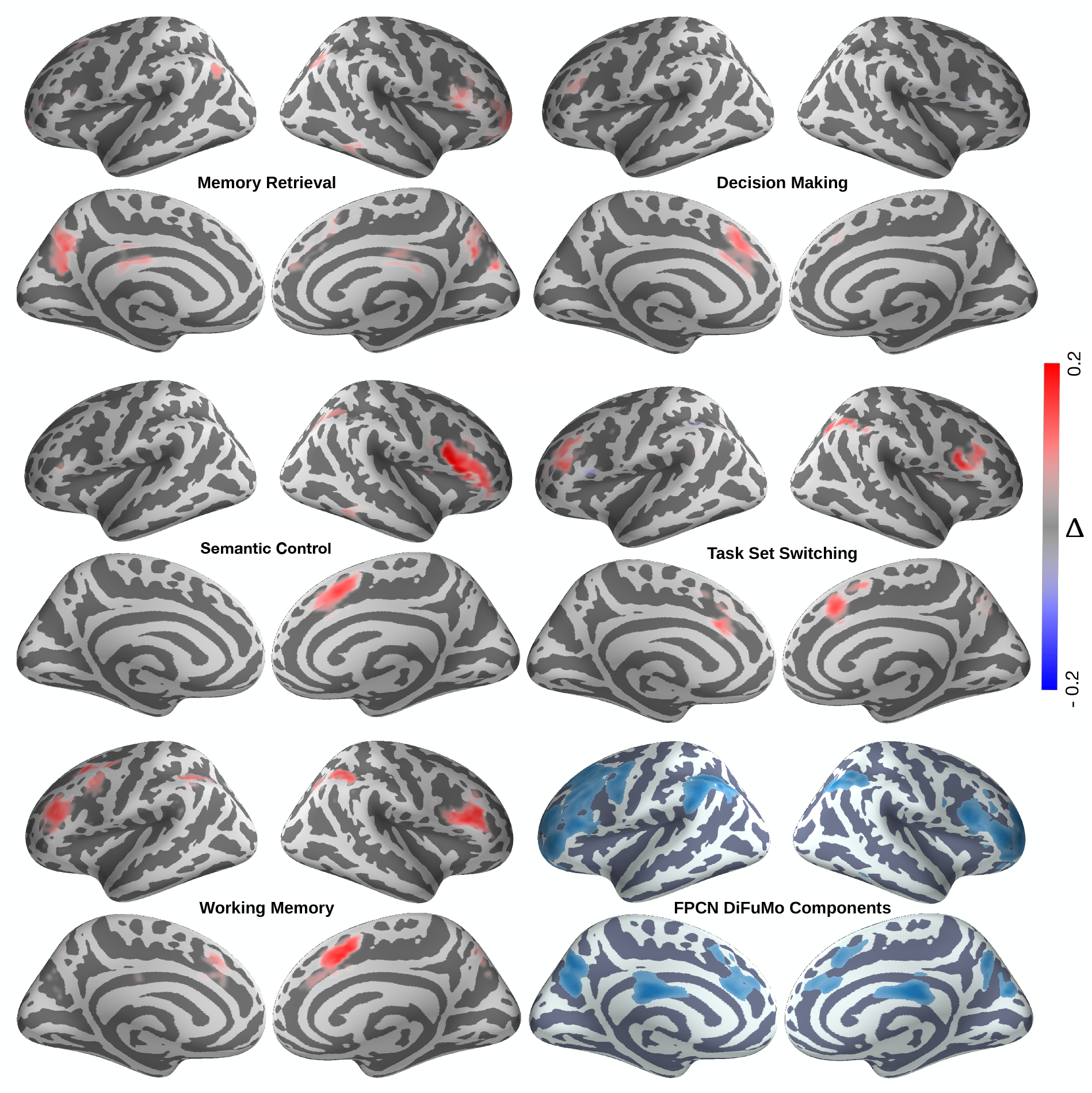
Cortical maps showing the difference in posterior probabilities of FPCN regions to be active given topic segregation and when given no topic segregation queries. We mask out brain voxels that are not part of the FPCN. The difference between posterior probabilities is defined as Δ = **P**[VoxelReported(*x, y, z*)|SingleTopicAssociation(*t*)] − **P**[VoxelReported(*x, y, z*)|TopicAssociation(*t*)]. Probability differences are thresholded at 0.05.

### Inferring Varying Meta-Analytic Connectivity Profiles of FPCN Subnetworks

Recent findings suggest that the frontoparietal cognitive control network (FPCN) can be decomposed into sub-systems associated with disparate and overlapping mental processes. Dixon et al. [7] studied two broad subsystems of the FPCN that also appear as separate networks in the influential 17-network model from Yeo et al. [21]. Using the same nomenclature, we label these two subsystems FPCN-A and FPCN-B. Dixon et al. observed preferential connectivity between FPCN-A and the default mode network (DMN), and between FPCN-B and the dorsal attention network (DAN). We reproduce these results by conducting a similar, but more compact, meta-analysis with NeuroLang. For this analysis we use the NeuroQuery [30] database instead of Neurosynth. We formulate conditional probabilistic queries that include studies reporting activations in each of the two FPCN sub-networks. By contrasting their posterior probability maps, we identify a distinct meta-analytic connectivity pattern associated with each sub-network. Using the same probabilistic definition of network reported by studies as in the first example, we formulate a rule that calculates the coactivation pattern of each FPCN subnetwork. In NeuroLang, we use the following rule to calculate the conditional probability of a region being reported given that a network is also reported

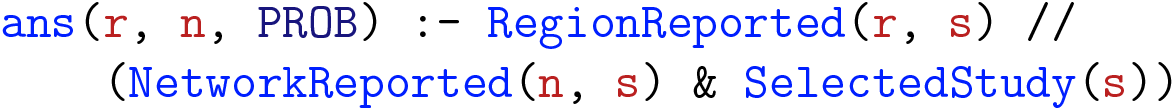

whose resulting ans table contains rows (*r, n, p*), where *p* is the probability of region *r* being reported active given that network *n*, where *n* is either FPCN-A or FPCN-B.

A likelihood-ratio test and a FDR correction (*α* = 0.05) for multiple comparison are used to identify statistically significant coactivating regions. To ensure that the results are not driven by one choice of studies, we estimate the conditional probabilities on 1000 random sub-samples of the NeuroQuery database (each sub-sample is 50% of the entire database). This allows us to compute confidence intervals for our probability estimates, which reflect the amount of variance across sub-samples.

In fig. 4, we show scatter plots of the probabilities that each DiFuMo-256 brain region is active given activation of the FPCN-A or FPCN-B sub-networks defined by Yeo 17-network parcellation. We only show the results of regions that exhibit a significant coactivation with at least one subnetwork, based on a likelihood-ratio test. In the left panel, regions are color coded by their network membership according to the coarser Yeo 7-network parcellation to facilitate interpretation.

**Figure 4:**
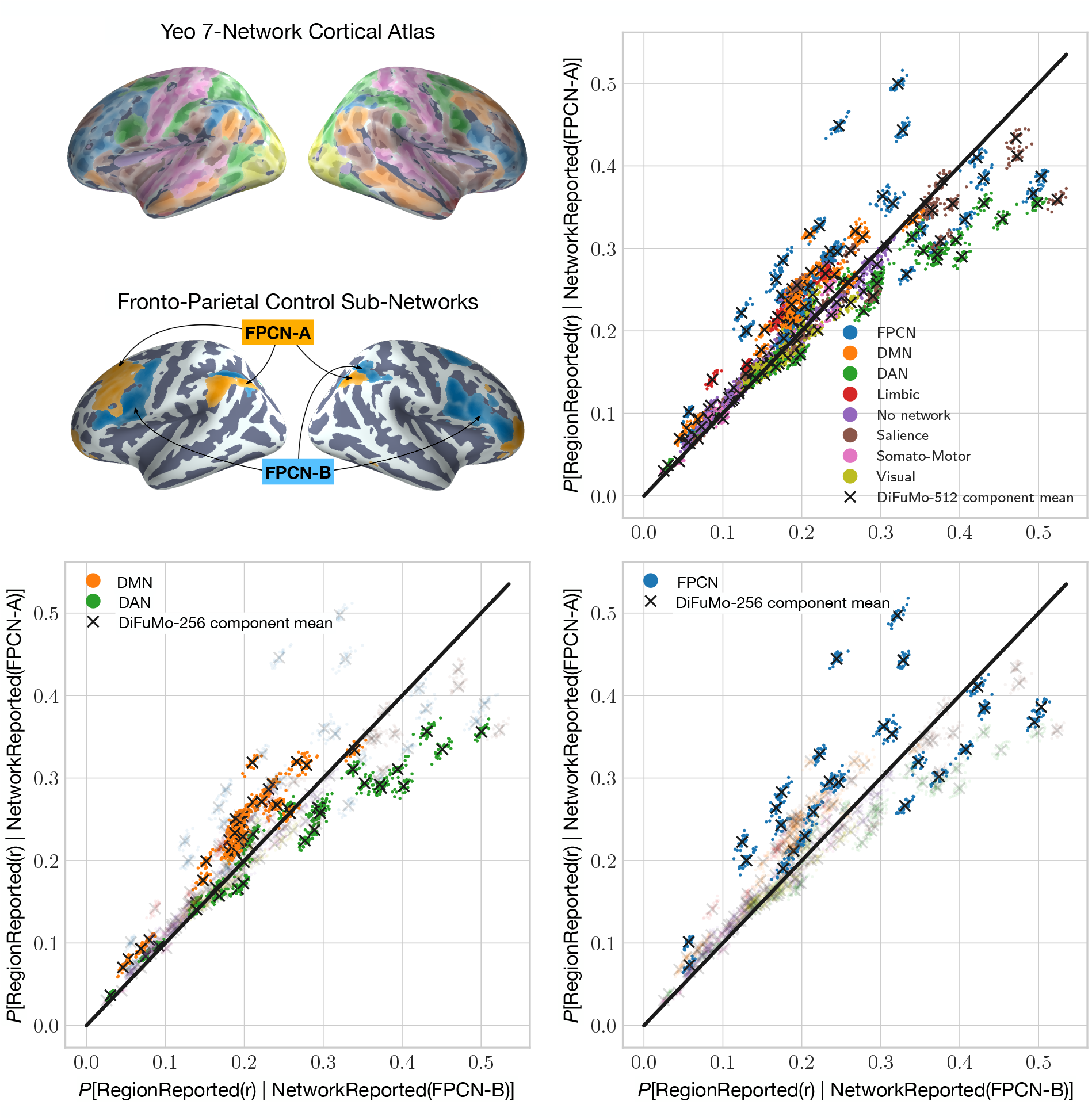
Comparison of the probabilities that DiFuMo-256 components coactivate with the two FPCN subnetworks. Regions are colored based on their belonging to the networks proposed by Yeo et al. [21]. Only regions exhibiting a statistically significant (*p*_FDR_ < 0.05) coactivations with either subnetwork are included in the figure, based on the likelihood-ratio test and a correction for multiple comparison. **P**[RegionReported(*r*)|NetworkReported(FPCN-A)] denotes the conditional probability of region *r* being reported by studies reporting FPCN-A in the database. Probabilities are calculated on 1000 random 50% subsamples of the NeuroQuery CBMA database.

In general, regions belonging to the somatomotor, visual, salience networks do not preferential coactivate with either the FPCN-A or FPCN-B. In contrast, regions of the coarse FPCN show a dichotomy in their coactivations with either FPCN-A and FPCN-B. That is, meta-analysis supports the hypothesis that FPCN can be functionally divided into two sub-systems [7]. Importantly, we find a clearer dichotomy in the coactivation profiles of the DMN and the DAN with the FPCN sub-networks. On one hand, 31 out of 32 DMN regions coactivate more with FPCN-A, while only one DMN region (a sub-region in the middle frontal gyrus) seem to exhibit a preferential coactivation with FPCN-B. In fig. 5, we illustrate a meta-analytic coactivation contrast map between FPCN-A and FPCN-B, showing that the former coactivates to a greater extent with the core regions of the DMN, than does the latter. On the other hand, without indicating any preference, we observe that 21 out of 30 DAN regions exhibit statistically significant coactivations with FPCN-A, while 19 show significant coactivations with FPCN-B. However, only 11 regions have a higher probability of activating given an FPCN-B activation than FPCN-A, while the others have comparable probabilities of coactivating with either sub-networks. This is in line with the findings from Dixon et al. [7], showing less distinction in the DAN with respect to coactivation with the FPCN sub-networks. Nonetheless, FPCN-B does indeed coactivate to a greater extent with the core regions of the DAN, the superior parietal lobule and frontal eye fields, than FPCN-A as seen from the coactivation contrast map in fig. 5.

**Figure 5:**
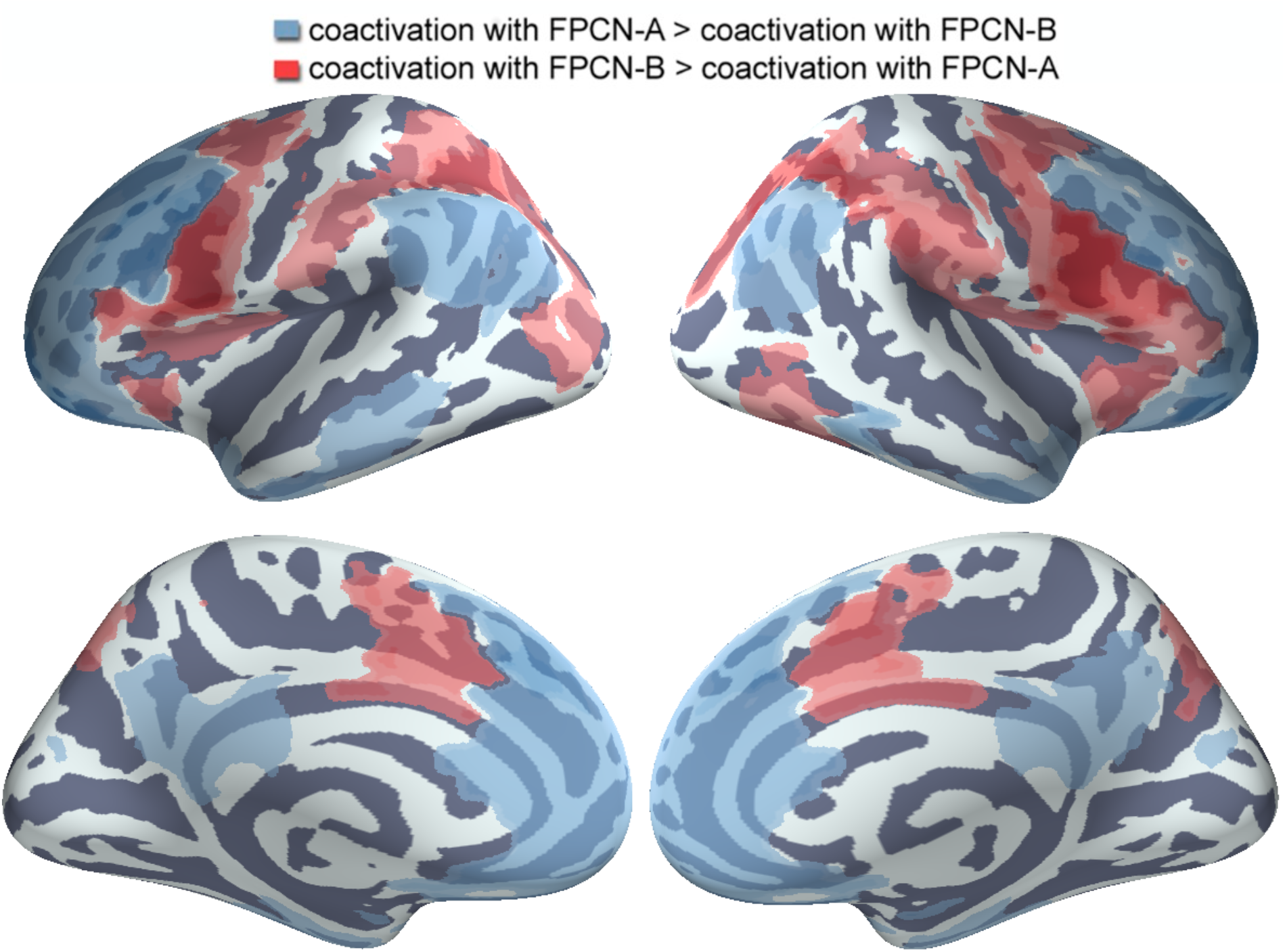
DiFuMo-256 components that are more likely to coactivate with one FPCN sub-network than the other. In blue, we depict regions exhibiting a greater probability of coactivation with FPCN-A. In red, we depict regions exhibiting a greater probability of coactivation with FPCN-B. The absolute difference between region coactivation probabilities is defined as Δ = **P**[RegionReported(*r*)|NetworkReported(FPCN-A)] − **P**[RegionReported(*r*)|NetworkReported(FPCN-B)]. A likelihood-ratio test and a FDR correction (*α* = 0.05) for multiple comparison are used to identify regions that exhibit significant coactivation with either networks before estimating Δ.

## 3 Discussion

We present a domain-specific language (DSL), coined NeuroLang, which broadens the range of meta-analytic hypotheses that can be expressed and tested against an ever-increasing functional neuroimaging literature. Through probabilistic logic semantics, users can formally represent their hypotheses, query heterogeneous data, and reason about uncertainty in a unified language. Ultimately, NeuroLang is envisioned to lead a new generation of computational tools for neuroimaging data analysis, including metaanalysis, to reduce miscommunication in the community and promote formal and reproducible research. To support this language-oriented approach, we provide concrete meta-analysis examples fully performed with NeuroLang.

An important meta-analysis application, beyond finding consistent activation patterns, is inferring reliable and *specific* structure-function associations. Traditionally, researchers carefully select studies that “ask a similar question” or rely on databases of expertly curated annotations of studies, such as BrainMap [2]. However, non-automated meta-analysis is not scalable, time-consuming, and can suffer from low statistical power. As a response, automated tools have been developed, like Neurosynth [3], NeuroQuery [30], and more recently NiMare, to enable scalable, richer, and unbiased meta-analysis. However, as mentioned before, these tools cannot formally express nontrivial inclusion/exclusion criteria of studies to infer specific structure-function associations. In contrast, NeuroLang provides a unified formal framework to succinctly express versatile queries of functional specificity in the brain via *first-order logic semantics* rather than propositional logic statements [19].

A recurring use-case of expressive querying throughout the examples is that of *segregation queries*, which we formulate using the first-order logical negation operator (¬) and existential quantifier (∃). Using a segregation query enables a seamless split-up of studies in a meta-analysis, while contrasting any number of topics and brain regions of interest both in a forward and reverse inference paradigms. In this sense, segregation queries can enhance the specificity of inferred structure-function associations in brain regions that are putatively recruited to varying degrees by multiple tasks and brain networks, such as the VWFA [24] or the anterior insula [31] for instance.

The three networks example serves as a proof of concept for using segregation queries to derive specific structure-function associations for brain structures, such as networks. In this example, we infer the topics that are preferentially linked to each of the FPCN, DMN, and DAN. These three networks are known to exhibit competitive and cooperative coupling dynamics across a wide array of tasks [32]. Thus, observed activations may emerge from the dynamics of these networks depending on task demands [33]. This might yield to a blurry characterization of networks in terms of their specific roles and spatial arrangement, sometimes leading to nomenclature ambiguities across studies and research groups [34]. One way to solve this is to infer relatively specific functions of a network by “isolating” its activity pattern from the other networks. Achieving this isolation through segregation queries, we find preferential topic associations for the FPCN, DMN, and DAN that align with the general understanding of their roles [26, 35, 36]. Such a segregation-based meta-analysis can be performed in future studies, for instance, to create a foundation for a fine-scale taxonomy of brain networks depending on their inferred roles in addition to their connectivity patterns.

In the VWFA example, we use segregation queries to address a controversial hypotheses [24, 37] about the role of this region in general visual processing beyond reading words. The main result is a statistically significant association for the VWFA with a topic loading on terms of object recognition given coactivation with attention but not language regions. However, the topic loads on terms such as ‘fusiform’ and ‘occipitotemporal’—zones in the close vicinity of the putative VWFA known to represent features and attributes of objects [38], which might bias the results due to uncertainty in precisely locating the VWFA and the ambiguity in nomenclature across studies. But given that the dorsal attention network is active and its strong functional connectivity to the left occipito-temporal zone [37], a link between the VWFA and “object recognition” is plausible. Concurrently, we observe no statistically significant topic associations for the VWFA when it coactivates with language but not attention regions. Yet, if we do not exclude studies reporting the coactivity among language and attention networks, a significant association with a topic related to “reading words” is observed. This finding suggests that the VWFA might not be specific to language per se, but rather have a broader role at the interface of language and visuospatial attention. Findings from [24] suggest that the VWFA acts as a gateway between attention and language networks, such that the former amplifies the representations of written words so they may be conveyed to the latter. Our findings suggest that the VWFA is a more general visual processor that may be recruited in other visual tasks in addition to reading. This segregation-based meta-analysis can be extended to understand the dynamic roles of flexible regions in the brain, such as connector hubs.

In the third and fourth examples, we derive varying coactivation patterns within the FPCN revealing its heterogeneous organization. The FPCN comprises regions that coactivate across diverse conditions. Given the lack of formal definitions and fine lines between different executive functions, which are often conjointly studied, it can be difficult to determine domain-specific FPCN regions. As a potential solution to this problem, a segregation query can simultaneously select studies highly loading on topic terms related to a single function, discard studies loading on topic terms of other functions, and contrast them. The results reveal a relatively unique coactivation pattern *consistently* associated with each topic, consistent with findings of dynamical activity in canonical brain networks as a function of varying demands [39]. We say “relatively unique” because we only study a small set of topics for the sake of demonstration, while in fact there are putatively more functions attributed to the FPCN. This meta-analysis can include a cognitive ontology [40] to systematically define *all* pertinent concepts and similarly contrast them [41]. Moreover, this type of meta-analysis can be effective in system-level causal modeling approaches (e.g., dynamic causal modeling) [42], which require strong a priori hypotheses about the regions involved in particular contexts. Finally, we reproduce the results of Dixon et al. [7], equivalently revealing two subsystems, FPCN-A and FPCN-B that exhibit distinct activations profiles and specific associations with the DMN and DAN, respectively. Although Dixon et al. [7] have successfully performed this analysis using Neurosynth’s command line tools, we have been able to reproduce their findings with significantly more compact, declarative and formal queries. In this sense, after representing the data in NeuroLang, a user only has to worry about what question to ask rather than about explicitly declaring every step needed to answer it.

Performing meta-analyses often requires integrating heterogeneous data. For example, Andrews et al. [11] use a brain parcellation to characterize components of the DMN whose respective functions are decoded through reverse inference reasoning. Here, to study the functional profiles of brain networks and the coactivation patterns of FPCN subsystems, we integrate the DiFuMo-256 functional atlas. Components of this functional atlas overlap with anatomical landmarks whose names are used to label the components by experts to enhance interpretability [20]. The DiFuMo components are also grouped into 7 and 17 canonical networks whose labels have been integrated in our examples. This approach facilitated the formulation of hypotheses and the interpretation of results. We believe that future studies conducted with NeuroLang could benefit from its capacity to represent any anatomical and functional atlas as well as tabular meta-data and formal ontologies [41]. Moreover, it is imperative for a *complete* meta-analytic tool to be flexible enough to represent any type of parcellation, meta-analytic database or, more generally, neuroscientific knowledge. Within our experiments, we have been able to represent both the Neurosynth database and its associated openly-shared topic models in NeuroLang. But, in other experiments, we use the NeuroQuery database because of its lower error-rate in the extraction of peak activation coordinates [30]. Together, these examples demonstrate that NeuroLang is agnostic to the database used for conducting meta-analyses, and could incorporate future sources of neuroscientific knowledge with various topologies.

Of course, due to the analytical variability across neuroimaging studies [43] and imperfections in data acquisition, knowledge representation should account for elements of uncertainty in the data. Probabilistic programs and databases constitute *general frameworks* for representing *structured* but *uncertain* knowledge. As these two paradigms reside at the heart of NeuroLang, uncertain data can be combined within its probabilistic programs. In our experiments, we model the reporting of functional modes and networks probabilistically based on their volumetric proportion that is reported by studies, although other indicators of uncertainty can be used. Another possible measure of uncertainty can be sample size and relative location of peaks with respect to the regions of a network. This probabilistic definition is thus *arbitrary*, but NeuroLang is expressive enough to represent any assumption just as well. For instance, to obtain network-specific functional profiles, we combine meta-analytic data and functional atlases with data-driven topic models. The topics are associated to studies probabilistically by the loadings of the fitted latent Dirichlet allocation model that are based on the frequency of co-occurrence of terms in abstracts of studies [18].

It is worth noting that, so far, not all types of queries can be *efficiently* solved by NeuroLang’s engine. For example, *negated* disjunctive queries, such as ¬(FirstSegregation(*s*) ∨ SecondSegregation(*s*)), where FirstSegregation and SecondSegregation correspond to two segregation rules — similar to those presented in our examples —, can be computationally intractable in a voxel-level analysis, where *>* 150, 000 voxels need to be modeled. This is because extensions to logic programming, such as negation [44], are not directly transposable to lifted query processing of unions of conjunctive queries on *probabilistic* databases [45]. In other words, more expressivity in a language leads to higher *data complexity* —the complexity of solving a query with respect to the size of the data [46]. In the case of whole-brain voxel-level statistical modeling, data complexity is significant, and solving a segregation query can become impractical due to the large number of voxels that are modeled. Efficient algorithms exist for solving a limited *subset* of queries on probabilistic databases [14]. NeuroLang privileges *lifted query processing* to solve queries on probabilistic databases (see the Methods section for details). Lifted query processing is an algorithm with a set of rules that translate a query to an algebraic expression to solve it in polynomial time. This allows NeuroLang to efficiently scale to large databases. However, to calculate the solution of a potentially non-liftable query, NeuroLang falls back to knowledge compilation strategies [47]. Yet, when modeling hundreds of thousands of brain voxels, we find resolution times of knowledge compilation to be impractical as well. Obtaining a solution for meta-analytic queries at the voxel-level currently take several minutes to be solved with NeuroLang, while such queries can be solved in a few seconds by Neurosynth’s engine, which uses a custom python-based implementation. Nevertheless, major improvements to NeuroLang’s engine are currently underway.

In designing NeuroLang, we provide a high-level programming interface for harnessing meta-analysis databases in cognitive neuroscience research. We believe that this approach has *three* main advantages : Accessibility, Readability, and Sound semantics. Implementing a program to test a complex hypothesis against meta-analytic data can be time-consuming and error-prone, especially for those not proficient in general-purpose programming languages. A domain-specific syntax as NeuroLang’s eases the process of formulating hypotheses combining heterogeneous data. This has the potential of speeding up the meta-analysis process as well as making it highly reproducible. Moreover, being specific about the research question and assumptions a in meta-analysis is an important practice [48]. NeuroLang’s logical syntax makes model assumptions and inclusion/exclusion criteria readable and understandable directly from the code of the program. Finally, NeuroLang is grounded in formal mathematical logic, providing theoretical guarantees that both limit modeling errors and provide trust in the language. We believe that these advantages make NeuroLang a tool of choice for conducting and reproducing functional neuroimaging meta-analyses. One NeuroLang program developed for a study in 2022 could be re-used by another team of researchers in 2027, five years later, either as a modeling inspiration for an entirely different study or to confront the original study’s findings with results published from 2022 to 2027.

## 4 Methods

We apply a language-oriented programming approach to the problem of expressing and testing meta-analytic neuroscientific hypotheses. That is, instead of using a general-purpose programming language to solve the problem, we design a domain-specific language (DSL) to represent the problem, and solve it in that language. NeuroLang uses *logic, declarative*, and *probabilistic* programming paradigms.

We start by introducing computer science concepts required to understand the semantics of the language. Then, we explore how these formalisms can be applied to the specific case of expressing cognitive neuroscience hypotheses. Finally, we detail the technicalities of solving queries on NeuroLang programs when working at the whole-brain neuroimaging scale.

### Probabilistic Logic Programs and Databases

To *represent* heterogeneous neuroscientific knowledge, NeuroLang leans on Datalog [49], a fully declarative logic programming language designed to efficiently solve queries on large deductive databases. A Datalog rule takes the form ∀**x**, (*ψ*(**x**) ← ∃**y**, *φ*(**x, y**)), where **x** is a set of universally quantified variables, **y** is a (possibly empty) set of existentially quantified variables, *ψ* is a relational symbol, and *φ*(**x, y**) is a conjunctive logic formula over **x** ∪ **y**. For readability, the implicit ∀ and ∃ quantifiers are often omitted, and the rule is written *ψ*(**x**) ← *φ*(**x, y**). A Datalog program is a set of such rules. The input of the program is a set of *extensional* facts and its output is a set of *intensional* facts that have been obtained through deductive inference, based on the program’s rules and input facts. This process is summarised in fig. 6. In NeuroLang, we write a Datalog rule ∀(*x, z*), (*P* (*x, z*) ← ∃*y, Q*(*x, y, z*) ∧ *R*(*z*)) as P(x, z) :Q(x, y, z) & R(z). Theoretical formalisms and efficient query resolution algorithms have been developed by the logic programming community over the past decades. By restricting the syntax of its rules, Datalog queries have been proven to be solvable in polynomial time w.r.t. the size of the database [49]. Strong guarantees on query resolution complexity are primordial to handle high dimensional whole-brain neuroimaging data.

**Figure 6:**
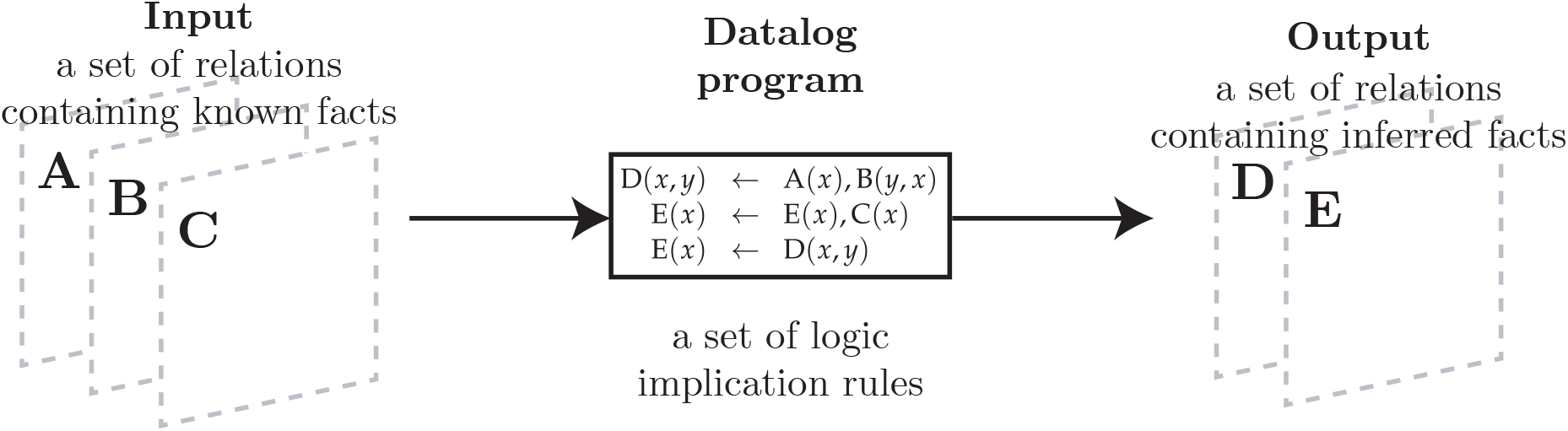
Deductive inference in Datalog. Using knowledge and implication rules to infer new knowledge.

**Figure 7:**
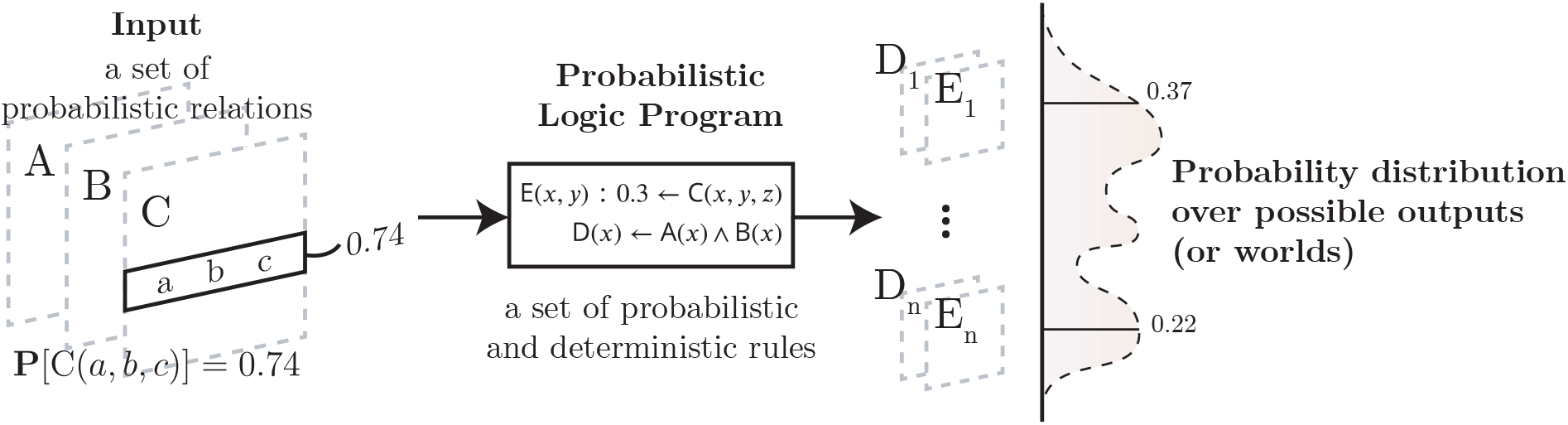
A probabilistic logic program defines a probability distribution over its possible outcomes.

Testing cognitive neuroscience hypotheses requires aggregating data from many subjects or studies using statistical models. Logic programming languages were not designed for statistics or probabilistic modeling, as their programs live in a world where everything is either true or false and only one outcome is possible. To address this limitation, logic programming languages were extended with probabilistic semantics to incorporate uncertainty and allow for probabilistic inference, such as ProbLog2 [50] or CP-Logic [51]. Following CP-Logic’s semantics, a probabilistic rule *ψ*(**x**) : *α* ← *φ*(**x**) causes the event *ψ*(**x**) to be true with probability *α*, whenever *φ*(**x**) is true. Probabilistic logic programs define a distribution over possible outputs of the programs’ execution, a.k.a. *possible worlds* [52]. For a more detailed introduction to probabilistic logic programming, we refer the reader to De Raedt et al.’s review [53]. In NeuroLang, we write a CP-Logic rule *P* (*x, y*) : *f* (*x, y*) ← *Q*(*x, y*) ∧ *R*(*y*) as P(x, y) : f(x, y) :-Q(x, y) & R(y).

Inferring the probability of a query boils down to summing the probabilities of all possible worlds where this query is verified. This process is called *weighted model counting* [54]. The number of possible worlds is often very large, and naively counting possible worlds becomes intractable. For example, were we to model the activation of each brain voxel as independent Bernoulli random variables, the number of possible worlds would be 2^*K*^, where *K* is the number of voxels in the brain, typically hundreds of thousands. Solving weighed model counting problems on real-world data requires efficient resolution algorithms. Knowledge compilation finds compact representations of large probabilistic programs, and can be used to solve queries drastically faster [14, 55].

In parallel, the field of probabilistic databases extended traditional relational databases with the possibility of encoding uncertain knowledge using probabilities [56]. A probability can be attached to any tuple in the database, as illustrated in fig. 8. Similarly to probabilistic programs that define a distribution over possible outputs, a probabilistic database defines a probability distribution over a set of possible databases (or worlds), where tuples are chosen to be true or false based on their probability. The probability attached to a tuple then corresponds to the marginal probability of that tuple being found in any database randomly chosen from the distribution over possible databases. Probabilistic tuples within the database are often assumed to be independent random events, in which case the database is called a tuple-independant database (TID). A tuple with probability 1 is true in all possible databases, and a tuple with probability 0.5 is true in half of the possible databases.

**Figure 8:**
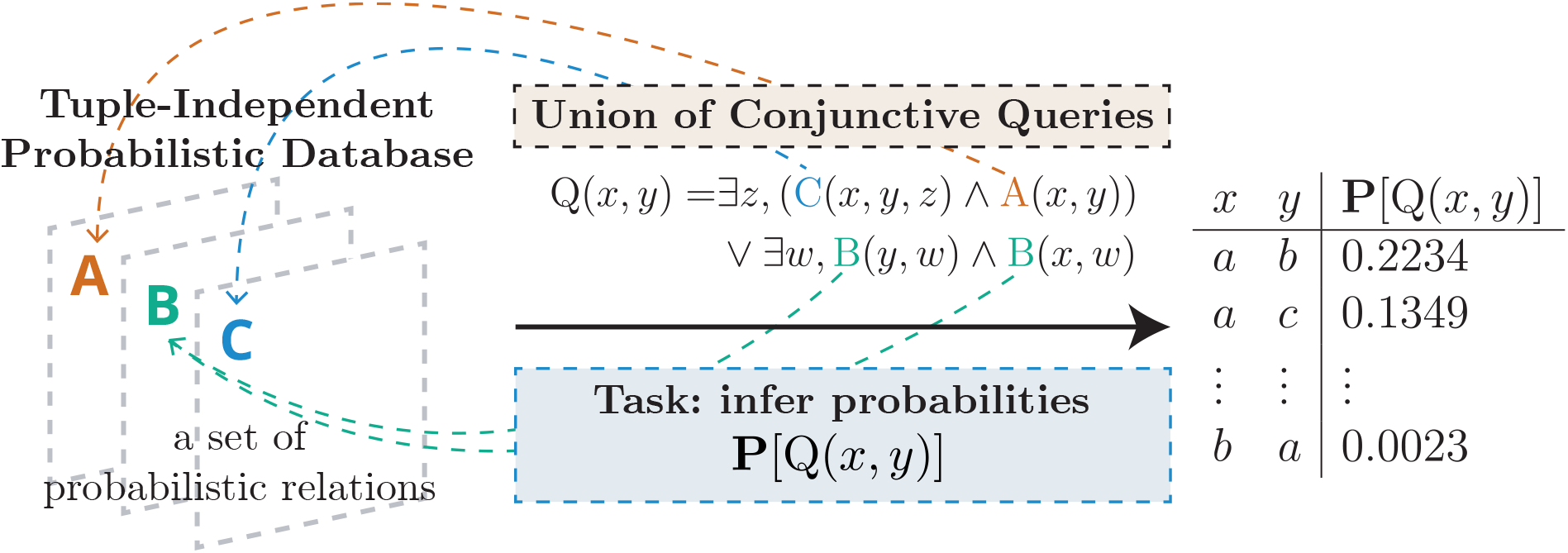
Solving a union of conjunctive queries on a tuple-independant database.

In some cases, solving a query on a probabilistic database can be done much more efficiently than through knowledge compilation approaches. One recent theoretical result is the *dichotomy theorem*, which classifies queries on TIDs based on their complexity [13]. At the heart of this theorem, is a resolution strategy named *lifted query processing*. It applies, based solely on syntactic analysis of a queries, a set of rules that derive an algebraic expression that computes the probability of the query [14]. We illustrate this process in fig. 9. Liftable queries have a polynomial data complexity. If the rules fail to apply, the query is said to be *non-liftable*, and has been proven to have a #P-hard complexity. To solve non-liftable queries, we use knowledge compilation strategies, which can be intractable in the voxel-level modeling setting.

**Figure 9:**
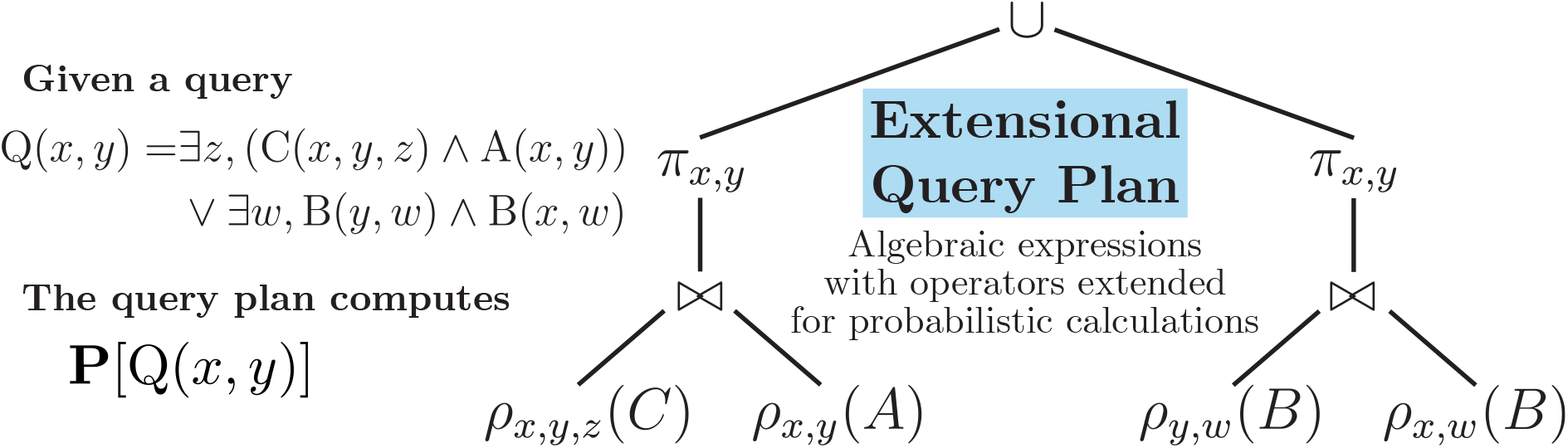
An extensional query plan solves **P**[*Q*] for a given query *Q* by using algebraic operations that are extended with probabilistic calculations [14].

By incorporating probabilistic semantics and fast query resolution algorithms from both probabilistic logic programming and probabilistic databases, NeuroLang is a fullfledged *probabilistic* programming language [19]. This approach makes it possible to express a wide variety of programs and queries, some of which can be efficiently solved using lifted query processing on probabilistic databases, even at the voxel-level.

### Syntactic Specificities of NeuroLang

We describe the syntactic extensions of typical logic and probabilistic logic programming languages that we made to provide features necessary to express end-to-end meta-analyses in NeuroLang.

A NeuroLang probability encoding rule (PER) captures the result of a probabilistic inference into a deterministic table. They are a syntactic convenience, or *sugar syntax*, that makes it possible to solve probabilistic queries within the program and process their solution with deterministic rules. Internally, we *stratify* (i.e. split in several code sections) a NeuroLang program as deterministic and probabilistic *strata*. Stratification allows us to *programmatically* post-process and analyse results from probabilistic calculations within the same self-contained program, as illustrated in fig. 10. Morever, this strategy can be used to supplement deterministic strata with logic programming extensions that would not necessarily be compatible with probabilistic programming [41].

**Figure 10:**
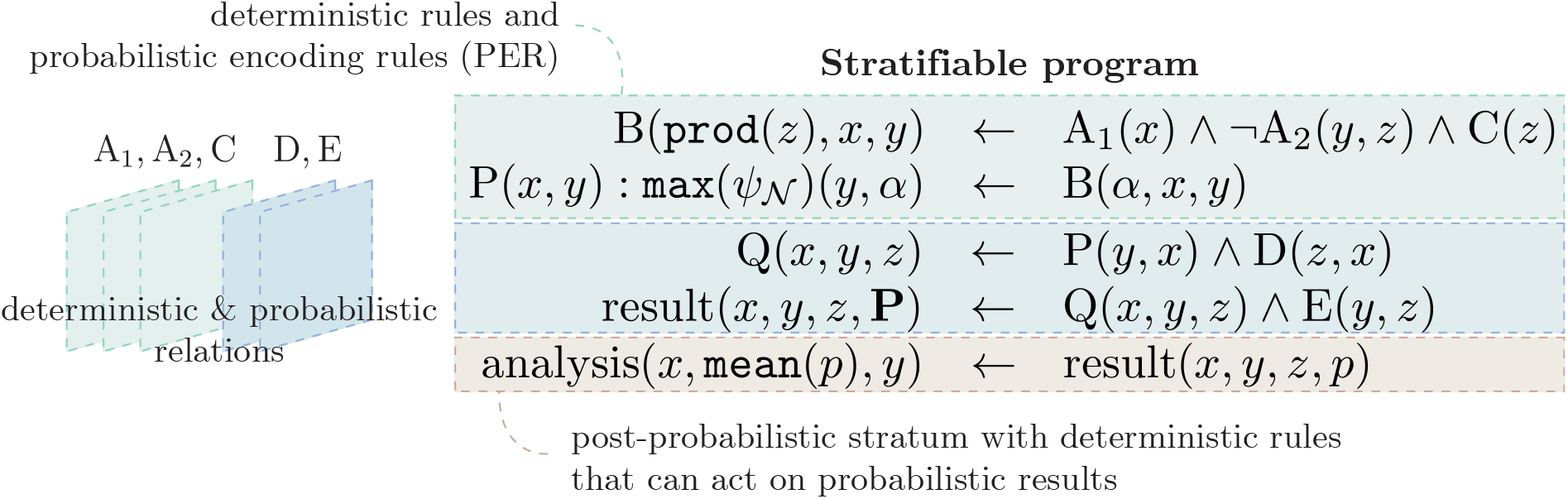
Stratification in NeuroLang. The program’s input contains both deterministic (A_1_, A_2_, and C), and probabilistic (D and E) tables.

PERs can either infer *marginal* or *conditional* probabilities. A marginal PER takes the form

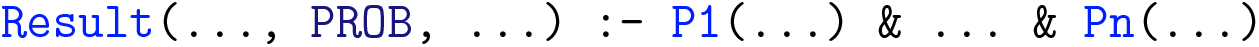

where P1, …, Pn are deterministic or probabilistic relational symbols, and where PROB is a special symbol that indicates which attribute of the resulting Result relation will contain the marginal probability that the rule’s antecedent is true. A conditional PER is only slightly different in that it calculates the probability of a conjunction of literals being true, *given that* another conjunction of literals is true. Conditional PER take the form

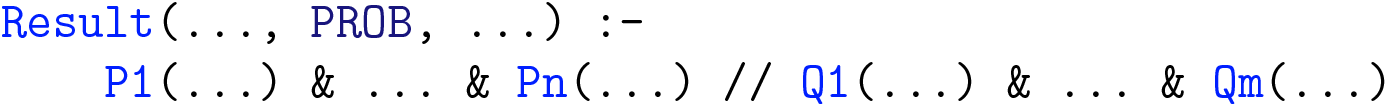

where the // operator applies a *probabilistic conditioning*. The PROB attribute of the resulting Result relation encodes the conditional probabilities **P**[P_1_(**x**)∧…∧P_*n*_(**x**)|Q_1_(**x**)∧ … ∧ Q_*m*_(**x**)], for each tuple **x** such that the probability is strictly positive.

NeuroLang supports explicit existential quantification of variables using rules of the form P(x) :-Q(x) & exists(y, R(x, y)), where variable y is existentially-quantified using the language’s special symbol exists. The language also supports negation within its deterministic and probabilistic rules, and aggregations within its deterministic rules. An example of aggregation rule is P(x, count(y, z)) :-Q(x, y, z), which counts, for each possible assignment of *x*, the number of tuples (*y, z*) such that *Q*(*x, y, z*) is true, and stores this count in the second column of table *P*.

Probabilistic tables can be constructed dynamically from deterministic rules. For example, the rule P(x) : f(x, y, z) :-Q(x, y, z) constructs a probabilistic table *P*, assigning a probability to each tuple (*x*,) based on function *f*. In the case where such rule generates multiple probabilities for the same tuple (*x*,), an error is thrown and the user is advised to either change her probabilistic definition, or apply an aggregation function on the probabilities, such as max(f(x, y, z)). The antecedent of these rules must be deterministic.

### Likelihood Ratio Test for NeuroLang Queries

Throughout this work, we formulate meta-analytic conditional probabilistic queries of the form **P**[*φ*(*s*) *ψ*(*s*)], where *φ*(*s*) and *ψ*(*s*) are first-order logic formulas describing study-specific probabilistic events of interest; such as whether a region / network is reported by study *s*, or whether *s* is associated with a topic related to a particular psychological concept. For brevity, we write **P**[*φ*(*s*)|*ψ*(*s*)] instead of **P**[*φ*(*s*) = ⊤|*ψ*(*s*) =⊤], where *φ*(*s*) and *ψ*(*s*) are modeled as Bernoulli random variables that have a probability of being true (⊤) or false (⊥) in any possible execution of the probabilistic logic program. The formula *ψ*(*s*) imposes conditions that select studies that will be included in a meta-analysis. To test the statistical dependence of *φ*(*s*) on *ψ*(*s*), we use a likelihood ratio test, whose null (*H*_0_) and alternative (*H*_1_) hypotheses are

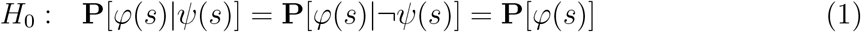

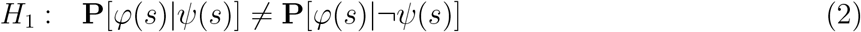

We define the likelihood ratio as *λ* = ℒ(*H*_1_)*/ℒ*(*H*_0_), where ℒ(*H*_1_) and ℒ(*H*_0_) are the maximum likelihood of the observed data under the alternative and null hypotheses, defined as

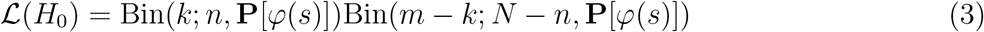

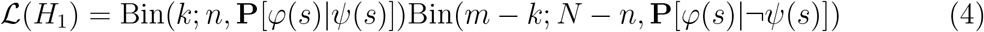

where *m* is the number of studies *s* such that *φ*(*s*), *n* is the number of studies *s* such that *ψ*(*s*), *k* is the number of studies *s* such that *φ*(*s*) ∧ *ψ*(*s*), and *N* is the total number of studies within the database. As 2 log *λ* is asymptotically *χ*^2^ distributed with 1 degree of freedom [57], it provides an estimate of the false-positive rate when rejecting the null hypothesis.

## 5 Acknowledgments

This work was funded by the ERC-2017-STG NeuroLang grant. We are grateful to Guillaume Favelier who helped with some of our visualisations. We are also grateful to Jonas Renault who worked on optimising NeuroLang’s engine and developing its webbased application. The source code of NeuroLang is openly available on GitHub at https://github.com/NeuroLang/NeuroLang.

