## Supplementary Material for "Meta-Analysis of the Functional Neuroimaging Literature with Probabilistic Logic Programming"

### SI Application of NeuroLang to the Resolution of Typical Queries on coordinate-based meta-analysis (CBMA) Databases

NeuroLang can be applied to the resolution of term-based and coactivation queries on CBMA databases, which are the most common types of meta-analyses conducted in the literature. A *term-based* query derives an activation pattern associated with a term of interest that relates to psychological concepts or cognitive process. A *coactivation* query delineates a brain activation pattern that comprises spatially distant brain regions, putatively forming a large-scale functional network. Answering these queries is typically done using tools like Neurosynth [1] or BrainMap [2].

#### SI.1 From Term to Association Map

A core goal of meta-analyses is to identify brain regions that are preferentially activated in experiments studying a psychological or cognitive process, relative to the rest of the brain [3]. As an example, a meta-analysis of studies on *emotion* should find that the amygdala is one of the regions that is the most likely to be reported by this set of studies. In other words, the meta-analysis should find that there is an association between studies of emotion and a neural response in the amygdala. Several models exists for computing probabilistic brain maps from a set of studies associated with a given term of interest ('emotion', in this example). All provide an estimate of the probability that a voxel gets reported by studies associated with the term. We explore and implement two of them in this example: multilevel kernel density analysis (MKDA) and activation likelihood estimation (ALE).

An important point should be made first, regarding the differences between approaches to combining the peak activation coordinates that are reported by neuroimaging studies. Peak activations are 3-dimensional points in the brain that live in a standardised stereotactic coordinate system, and that can be seen as a sparse representation of the statistical map obtained from an experimental study of neuroimaging signals. To account for the noise and variability of the *exact location* of these peaks, meta-analysis models often assume a neighborhood around peaks to be reported, or to have a probability of being reported. The definition of this spatial neighborhood is an *a priori* of the meta-analysis, sometimes called a spatial prior. This process, applied to the map of each study separately, is called *smoothing* [4].

A simple but common spatial smoothing approach is to consider all voxels within a sphere centered at a reported peak location's coordinates to also be reported by the study. This prior, used by MKDA, is illustrated on the left part of fig. S1. MKDA obtains one such binary pattern for each study, and aggregates

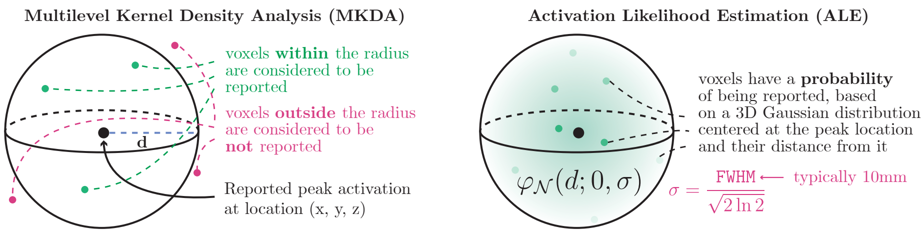


Figure S1: Illustration of spatial smoothing priors used by MKDA and ALE to address the uncertainty in the location of peak activation coordinates reported by neuroimaging studies.

them as frequencies at which each voxel is reported by studies selected for a meta-analysis. In plain English, 'a voxel at location $(x, y, z)$ is reported by a study $s$ whenever $s$ reports a peak activation within 10mm of that voxel'. In NeuroLang, this sentence is encoded by the following `VoxelReported` rule

```
VoxelReported(x, y, z, s) :-
    PeakReported(x2, y2, z2, s) & Voxel(x, y, z)
    & d = euclidean_distance(x, y, z, x2, y2, z2) & d < 10
```

where the pairwise Euclidean distance between pairs of voxels is calculated using a built-in `euclidean_distance` function. The lower bound `d < 10` defines a 10mm radius ball around each peak location [5]. An MKDA meta-analytic map is obtained by estimating the conditional probability that each voxel gets reported by studies associated with a given term of interest (e.g. 'emotion'.) To compute estimates of these probabilities, we define the following NeuroLang rule, which gives each study an equal probability of being selected

```
SelectedStudy(s_1) : 1/N, ..., SelectedStudy(s_N) : 1/N :- ⊤
```

The distribution of the program's outputs can then be used to estimate the probability of each voxel being reported. Then, the result of solving a conditional probabilistic rule

```
ans(x, y, z, PROB) :-
    VoxelReported(x, y, z, s) & SelectedStudy(s)
    // TermAssociation(emotion, s) & SelectedStudy(s)
```

will contain a probability for each voxel of the brain, based on the frequency at which it is reported by studies within the database. The `//` operator represents a *probabilistic conditioning*.

Another approach to smoothing peak activations is to model their location's uncertainty *probabilistically*. We give the example of the ALE algorithm [6], illustrated on the right part of fig. S1. Instead of using a hard sphere, a three-dimensional Gaussian probability distribution is defined at each peak. This distribution provides a probability that depends on the Euclidean distance between each reported peak and brain voxel. The standard deviation of the Gaussian distribution is directly linked to the user-specified full-width half-maximum (FWHM), as in the ALE algorithm. In NeuroLang, this is modeled using a rule that programmatically defines a `VoxelReported` table as follows

```
VoxelReported(x, y, z, s) : p :-
    PeakReported(x2, y2, z2, s) & Voxel(x, y, z)
    & d = euclidean_distance(x, y, z, x2, y2, z2) & d < 2 * FWHM
    & p = gaussian(d ; 0, sigma) & FWHM = 10
    & sigma = FWHM / 2 * sqrt(2 * log(2))
```

Following the ALE model, the probability of activation for voxel $(x, y, z)$ is calculated as the probability that any study in the meta-analysis report its activation. Assuming voxel reporting events to be independent, this means that the probability $\mathbf{P}[\exists s, \text{VoxelReported}(x, y, z, s) \wedge \text{TermAssociation}(\texttt{emotion}, s)]$ is equal to

$$1 - \prod_s (1 - \mathbf{P}[\text{VoxelReported}(x, y, z, s) \wedge \text{TermAssociation}(\texttt{emotion}, s)]) \quad (1)$$

where $s$ is a study, as described in [7]. To estimate this probability, we use an *existentially-quantified* rule — that is, a rule that uses a predicate logic existential ($\exists$) quantifier — as follows

```
ans(x, y, z, PROB) :- exists(s;
    VoxelReported(x, y, z, s)
    & TermAssociation("emotion", s)
)
```

In fig. S2, we compare the probability maps obtained from the MKDA and ALE NeuroLang programs. As expected, the association maps obtained with the two methods highlight a cluster in the amygdala, which is known to be part of the neural signature of emotion.

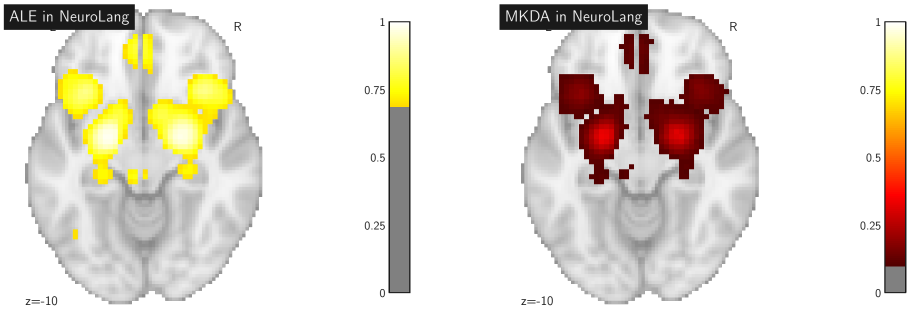


Figure S2: Probability maps for the term 'emotion', obtained from the MKDA and ALE NeuroLang programs. Values are shown before any statistical test is performed. The maps are thresholded to select only the top 1% probabilities.

### SI.2 From Seed-Voxel to Coactivation Map

We now show how meta-analyses of voxel coactivations can be formulated in NeuroLang. These analyses are able to capture some of the most studied functional networks of the brain: the frontoparietal cognitive control network, dorsal attention network and sensory-motor network. This was already shown to be possible on the manually curated BrainMap database [8]. We replicate these results, with NeuroLang, on the automatically extracted NeuroQuery CBMA database [9]. To solve coactivation queries in NeuroLang, we use the same program as in the previously described MKDA example, although the `TermAssociation` table plays no role now, as it is not needed for solving coactivation queries. We formulate the following conditional probabilistic rule

```
ans(x, y, z, PROB) :-
    VoxelReported(x, y, z, s) & SelectedStudy(s)
    // VoxelReported(x2, y2, z2, s) & SeedVoxel(x2, y2, z2)
      & SelectedStudy(s)
```

which asks, in plain English, 'what is the probability that each voxel at MNI coordinates $(x, y, z)$ gets reported by studies reporting the activation of some seed voxel at MNI coordinates $(x_2, y_2, z_2)$'. This is solved for every voxel within 10mm of any peak reported at least once in the database, because the MKDA spatial smoothing prior is used. Note that this rule computes the coactivation maps for each seed voxel within the `SeedVoxel` table, meaning that the query needs to be solved only once to obtain several meta-analytical results. The `SeedVoxel` table contains the same coordinates as the ones used by Toro et al. [8].

In fig. S3, we show the resulting coactivation map for each seed voxel. The first coactivation map, with a seed voxel in the left central sulcus (CS), correctly highlights the sensory-motor network (SMN), with clear activation clusters in the precentral and postcentral gyri [10]. The second coactivation map, with a seed voxel in the anterior cingulate cortex (aCC), correctly highlights the default mode network (DMN), with activation clusters in the medial prefrontal cortex, posterior cingulate cortex, and nucleus accumbens. Finally, the third coactivation map, with a seed voxel in the left intraparietal sulcus (lIPS), correctly hightlights the dorsal attention network (DAN), with activation clusters in the supplementary motor area, anterior insula, frontal eye fields, dorsolateral prefrontal cortex, and ventral occipital cortex. These coactivation maps correctly highlight the functional network their respective seed voxel is expected to belong to, similarly to what was reported by Toro et al. [8].

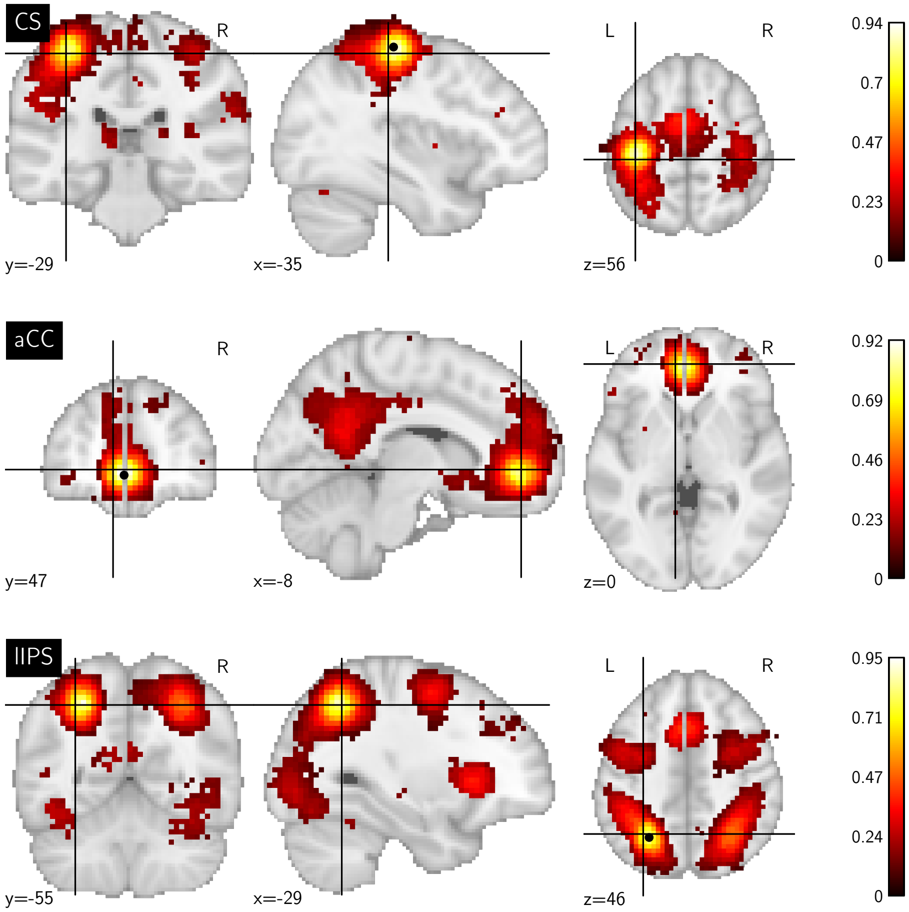


Figure S3: Coactivation maps obtained from seed voxels in the dorsal part of the left CS ($x = -34, y = -26, z = 60$), the aCC ($x = -2, y = 46, z = -4$), and the lIPS ($x = -26, y = -58, z = 48$). The number of studies reporting activations within 10mm of each seed voxel was 971, 1812, and 1380 1380, 1812 and 971 for the CS, the aCC, and the lIPS (respectively).

To conclude, we have shown that typical term-based and coactivation-based

| **Term associated with studies reporting the VWFA and the 'language' network, but not reporting the fronto-parietal attention network** | |
|---|---|
| Term | $p$-val (uncorrected) |
| l2 | 0.000773 |
| bilinguals | 0.001411 |
| l1 | 0.003706 |
| sublexical | 0.006793 |
| visual word recognition | 0.006987 |
| **Term associated with studies reporting the VWFA and the fronto-parietal attention network, but not reporting the 'language' network** | |
| Term | $p$-val (uncorrected) |
| occipitotemporal cortex | 0.000008 |
| object recognition | 0.000145 |
| conflict detection | 0.007809 |
| spatial processing | 0.007809 |
| visual working memory | 0.009181 |
| visual word recognition | 0.009684 |
| word recognition | 0.009684 |
| ba37 | 0.009684 |
| visual stream | 0.014894 |
| exner | 0.024605 |
| fonts | 0.032104 |

Table S1: Term associations surviving a likelihood ratio test ($p < 0.05$).

queries on CBMA could be expressed and solved with NeuroLang.

### SII VWFA: Term Associations not Surviving Correction for Multiple Comparison

In table S1, we complement our topic-based study of the visual word-form area (VWFA) but with an analysis of raw terms within the Neurosynth database. The terms are indeed related to the topics found in our experiments: object and word recognition. However, none of these term associations survive a FDR-based correction for multiple comparison. The calculation is ran for all terms within Neurosynth's vocabulary. We conducted the same analysis on the NeuroQuery database, which contains a broader vocabulary and more term associations, and obtained similar results.
